# Exclusion and genomic relatedness methods for assignment of parentage using genotyping-by-sequencing data

**DOI:** 10.1101/582585

**Authors:** K. G. Dodds, J. C. McEwan, R. Brauning, T. C. van Stijn, S. J. Rowe, K. M. McEwan, S. M. Clarke

**Author notes:** **Address for Correspondence** K. G. Dodds, AgResearch, Invermay Agricultural Centre, Private Bag 50034, Mosgiel 9053, New Zealand.

## Abstract

Genotypes are often used to assign parentage in agricultural and ecological settings. Sequencing can be used to obtain genotypes but does not provide unambiguous genotype calls, especially when sequencing depth is low in order to reduce costs. In that case, standard parentage analysis methods no longer apply. A strategy for using low-depth sequencing data for parentage assignment is developed here. It entails the use of relatedness estimates along with a metric termed excess mismatch rate which, for parent-offspring pairs or trios, is the difference between the observed mismatch rate and the rate expected under a model of inheritance and allele reads without error. When more than one putative parent has similar statistics, bootstrapping can provide a measure of the relatedness similarity. Putative parent-offspring trios can be further checked for consistency by comparing the offspring’s estimated inbreeding to half the parent relatedness. Suitable thresholds are required for each metric. These methods were applied to a deer breeding operation consisting of two herds of different breeds. Relatedness estimates were more in line with expectation when the herds were analysed separately than when combined, although this did not alter which parents were the best matches with each offspring. Parentage results were largely consistent with those based on a microsatellite parentage panel with three discordant parent assignments out of 1561. Two models are investigated to allow the parentage metrics to be calculated with non-random selection of alleles. The tools and strategies given here allow parentage to be assigned from low-depth sequencing data.

## Introduction

As the cost of sequencing declines, it becomes feasible to use this technology to obtain genomic information for research or commercial applications. Often sufficient information is given by sequencing a proportion of the genome, for example by using reduced representational sequencing approaches such as genotyping-by-sequencing (GBS; Elshire *et al.* (2011)), restriction-site associated DNA sequencing (RAD-seq; Baird *et al.* (2008)) or exon capture sequencing (Ng *et al.* 2009). To reduce costs further, sequencing may be undertaken at low depth. However this increases the chance of not reading both alleles at a locus, which may result in a heterozygous individual being called as homozygous for one of the alleles. An attractive feature of sequencing-based genotyping is that it does not require up-front costs of developing marker panels; SNPs are discovered and genotyped in the same process and this can be done in the absence of a reference genome sequence for the species.

A common use of genotype data is to assign parentage, for example in agricultural or ecological settings (Städele & Vigilant 2016; Grashei *et al.* 2018). Sequencing may provide a cost-competitive option for this task, especially if other genomic information (such as breed assignment or population structure) is also sought. Checking recorded pedigrees is useful for quality control of sample assignments. The assignment of dams allows the inference of non-genetic effects such as birth date and litter size (Dodds *et al.* 2005) and allows maternal models to be fitted, even if the relationships are estimated genomically. Similarly, if genomic prediction combines information from genotyped and ungenotyped individuals in a ‘single step’ analysis (Aguilar *et al.* 2010), some tuning parameters (Legarra *et al.* 2014) rely on there being pedigree information on all individuals.

As low-depth sequencing does not provide unambiguous genotype calls, standard parentage analysis methods no longer apply. A common approach to analysing such data is to filter out the genotype calls with low read depths (Kim *et al.* 2016). However, this greatly increases the proportion of missing data and may result in low numbers of SNPs called in both (all) members of a parent-offspring pair (trio). An approach that is used if parentage analysis is the primary objective is to filter to a set of high quality (including high call rate) SNPs (Thrasher *et al.* 2018). In both approaches, there is a loss of information in the discarded data. Here we investigate methods for parentage assignment from sequence data that take into account the way in which the genomic information is obtained, i.e. using a model of random allele reads at a particular SNP.

The methods developed here can be applied to parentage verification (where parentage has been recorded, but genotyping is used to check the parentage) or for parentage assignment (where parents are not recorded, but are known to come from a given group and genotyping is used to match to the specific parent(s)). This article will focus on the latter situation as it is more general.

## Methods

### Use of mismatch rates for parentage assignment

A common approach to parentage assignment is by exclusion: find individuals in the father and mother sets (ideally only one in each set) which have genotypes consistent with parentage, i.e. no “mismatches”. As (low-depth) GBS data does not always give the true genotype (heterozygous individuals are sometimes observed as homozygotes), there will be mismatches, even with the true parents, with non-zero probability. Here we consider mismatch rates (number of mismatches divided by the number of comparisons) as these will be more stable over differing numbers of comparisons than mismatch counts. One possible approach is to calculate the observed mismatch rate after filtering the genotype calls to a given minimum depth. However, this will reduce the data available for a comparison, especially when trios are considered. Instead we calculate the “excess mismatch rate” (EMM) as the observed mismatch rate minus the expected mismatch rate.

The expected mismatch rate is calculated under the hypothesis of parentage and takes into account the read depths of the individual and the putative parent(s). The derivation is given using the model (and some possible extensions) and notation of Dodds *et al.* (2015). Let A and B denote the alleles of a SNP and *g*\* denote the apparent genotype (e.g. AA* denotes that only A alleles are observed). Suppose homozygotes are observed without error, but for heterozygotes

*P*(AA*|AB) = *P*(BB*|AB) = *K* and

*P*(AB*|AB) = 1 − 2*K*

The binomial model assumes that allele reads are at random, leading to *K* = 1/2^*k*^ for a genotype with depth *k*. The theory is presented in terms of *K* for simplicity of presentation and to allow other sampling models (relating *K* to *k*) to be easily implemented.

Expected mismatch rates are calculated under the assumption of a randomly mating, non-inbred population in Hardy-Weinberg equilibrium. Let *p* be the A allele frequency (assumed known). An example of probability calculations for a set of true and observed genotypes is given in Table 1. The full table is given in Table S1. The corresponding probabilities for a single parent and offspring genotypes is given in Table S2 and the derivation of the probability of an apparent mismatch, given the offspring apparent genotype, read depth and read depths of the parent(s) is given in the Supplemental material.

**Table 1.** Example of probability calculations for the case where the true genotypes for father, mother and offspring are AA, AB and AA, respectively. *K*_*m*_ is the value of *K* for the mother.

| Observed genotype |  |  | Probability | Observed Mismatch? |
| --- | --- | --- | --- | --- |
| Father | Mother | Offspring |  |  |
| AA | AA | AA | $p^3(1-p)K_m$ | |
| AA | AB | AA | $p^3(1-p)(1-2K_m)$ | |
| AA | BB | AA | $p^3(1-p)K_m$ | Yes |

Let *K*_*x*_ with *x* = *o,m,f* to denote the value of *K* for the offspring, putative mother and putative father, respectively. The probabilities of an apparent parent-pair-offspring mismatch, given the offspring genotype, are

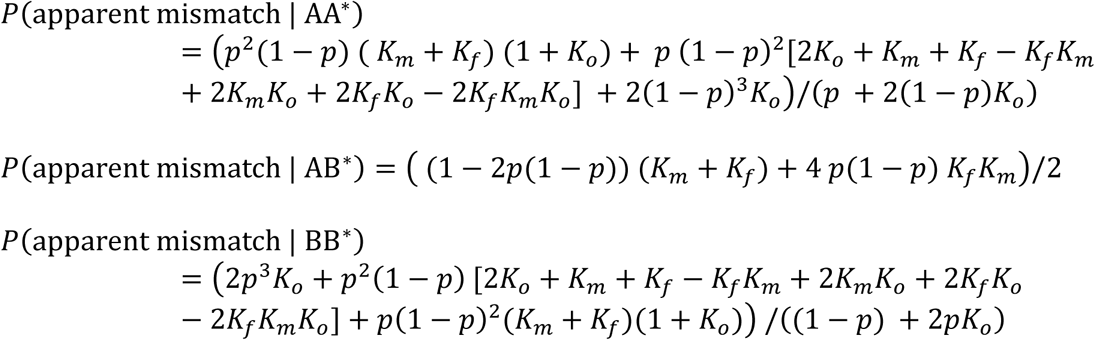

The probabilities of an apparent father-offspring mismatch, given the offspring genotype, are

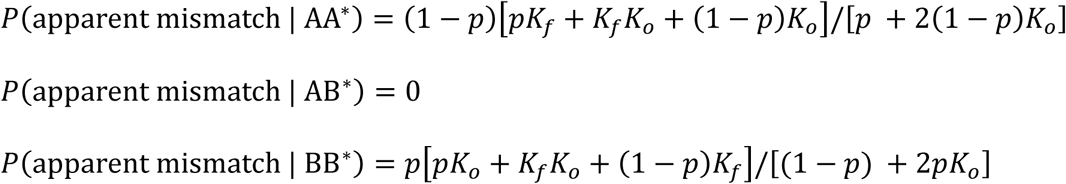

The equations for mother-offspring mismatch are the same, but with *K*_*f*_ replaced by *K*_*m*_. The expected mismatch rate for an offspring is calculated as the average of the relevant set of these equations over all the SNPs.

### Use of relatedness estimates to assign parentage

We consider an approach (Moore *et al.* 2019) that uses a relatedness estimator appropriate for the analysis of GBS data (Dodds *et al.* 2015). Let *r* and *F* denote relatedness and inbreeding, respectively, and subscripts O, F, and M denote the offspring, father and mother, respectively. For non-inbred offspring and parents, *r*_OF_ = *r*_OM_ = 0.5. Allowing for related and possibly inbred parents, we have, e.g., *r*_OF_ = 0.5 + 0.5*F*_F_ + *F*_O_ (Grashei *et al.* 2018); the relatedness can be greater than 0.5.

Genomic-based estimates give relatedness relative to some (often not well defined) base population (Powell *et al.* 2010), and so even a non-inbred individual and a parent may not have a relatedness estimate close to 0.5. A possible solution is to find a transformation (Weir & Goudet 2017) so that the parents and their non-inbred offspring have estimates near 0.5, for example by using pairs of known (pedigree-based) relatedness or scaling so that minimum estimates are close to zero (if it is reasonable to assume some members of the population are unrelated). In many situations, there will be limited sets of possible male and female parents, and the true parents will be the member of each set that has the highest relatedness with the offspring in question. Any reference to relatedness or inbreeding in the following will be taken to mean estimated values.

### Parentage assignment strategy

A strategy for parentage assignment, based primarily on either EMM or relatedness metrics, follows. Here “best” is taken to mean either the lowest EMM or the highest related, accordingly.

1) Initially assign the best member of possible fathers as the father
2) Initially assign the best member of possible mothers as the mother
3) Discard any assignments that fall beyond some threshold (EMM too high or relatedness too low), for example by visually examining the distribution of metrics for the initial assignments.
4) Discard any initial assignments where the metrics for the best and second best member of the possible parents are deemed too close to accept the initial assignment.
5) Check that the combined (trio) assignment is consistent (both the trio EMM and *r*_*FM*_ − 2*F*_*O*_ are not too high).

If only one parent gender is being assigned (and the other parent is unknown or not genotyped), then any of the steps above that involve using both parent assignments together will not be relevant. Even though one of the methods (EMM or relatedness) must be chosen to find the best parent, thresholds for both methods can be used jointly to discard dubious assignments.

The methods presented here have been incorporated into R code available at https://github.com/AgResearch/KGD. This code is more efficient at calculating relatedness than EMM, and so the relatedness approach would be preferable if the performance of the two approaches is otherwise similar. For the relatedness approach, a bootstrapping method is investigated to aid step 4. The SNP markers are sampled with replacement to obtain a bootstrap sample with the same number of SNPs and the relatedness values recalculated. The proportion of times across many (e.g. 1000) bootstrap samples that the most related parent has a higher estimated relatedness (with the offspring) than the second most related parent is denoted as the bootstrap support for the assignment.

In step 5 when using relatedness, a low value of *r*_*FM*_ − 2*F*_*O*_ might also be rejected. In practice, it should suffice to place an upper limit on this value, as the likely errors are when one parent is correctly assigned, and the other parent is incorrectly assigned, due to its high relatedness with the correctly assigned parent.

### Alternative allele sampling models

There may be models that are better for modelling the data than the binomial model, for example ones where allele reads exhibit clustering such that observed homozygosity is higher (when depth exceeds one) than in the random reads case. The calculations here require *K*, the probability of all A alleles (no B alleles) for a true AB genotype, and only consider models where this is the same as the probability of all B alleles. In particular, *K* = ½ when *k* = 1.

One possible alternative model is the beta-binomial (BB) model, which is often used to model data that are more dispersed than a binomial, for example, where there is some hidden clustering of the sampling process. The probability of seeing *m* B alleles with depth *k* is

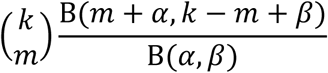

where B is the beta function. The mean of the distribution, with *k* = 1, is α /(α + β) which is equal to ½ when β = α. Making this substitution and setting *m* = 0,

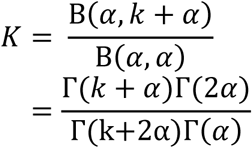

where Γ denotes the gamma function. This model approaches the binomial model as α approaches infinity.

The second alterative model considered assumes that the sampling process for reads for a particular SNP is Markovian – the probability of reading a particular allele depends on which allele was seen for the previous read (of that SNP). We denote the probability of seeing the same allele as was previously read as *p*′. The binomial model is a special case with *p*′=0.5. We refer to this as the ‘modified *p*’ (MP) model. As noted above, the probability of seeing a particular allele for the first read (the only read if *k*=1) is 0.5. Because we are only interested in the cases where all the reads are the same, we do not actually need to know the order that the alleles were read. For this model

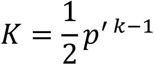

Examples of these models are shown in Figure S1. Both these models need the value of an extra parameter. Here we estimate the parameter by finding a set of parent-progeny trios that are assumed correct based on the binomial model (which is more conservative for calculating expected mismatch rates). The sum of squares of the deviations of the mismatch rates from the expected mismatch rates for these trios is then minimised (using the *optimize* function in R) with respect to the unknown parameter.

The alternative sampling models will alter the expected mismatch rates and the self-relatedness (1+*F*) estimates, but not the relatedness estimates between individuals, as these do not depend on *K* (Dodds *et al.* 2015).

### Animals

The methods have been applied in two herds of deer, part of Focus Genetics’ breeding programme (www.focusgenetics.com). One was a Red deer (*Cervus elaphus*) breeding herd and the other was a Wapiti (also known as Elk; *Cervus canadensis*) breeding herd. The animals were managed in accordance with the provisions of the New Zealand Animal Welfare Act 1999, and the Codes of Welfare developed under sections 68–79 of the Act. Even though Red deer and Elk are considered different species (Pitra *et al.* 2004), they are capable of inter-breeding and produce fertile progeny. We will refer to them as breeds here, as that is how they are considered in New Zealand deer farming. Although the herds are not necessarily purebred, they are denoted here by their predominant breed.

There were 1272 Red deer successfully genotyped by GBS (mean read depth of at least 0.3 over all SNPs called). These consisted of 571 progeny, born in 2015, 34 potential sires (none missing) and 667 potential dams (10 missing, i.e., unavailable or not successfully genotyped). There were 709 Wapiti successfully genotyped by GBS consisting of 246 progeny, born in 2015, 41 potential sires (none missing) and 422 potential dams (two missing). There may be additional sires and dams that were not sampled. This resource had also been genotyped using a panel of 16 microsatellite markers, a commercial in-house deer parentage panel used by GenomNZ (www.genomnz.co.nz) and parentage assignments made based on those genotypes.

### GBS Genotyping

Genotyping was undertaken based on the method of Elshire *et al.* (2011) as described in Dodds *et al.* (2015) apart from the following minor differences. DNA extraction was from ear-punch tissue. The GBS-libraries were prepared utilising the *Pst*I restriction enzyme. Either 96 or 192 samples were run per lane on an Illumina HiSeq2500 to obtain 100bp single end reads. The genotypes were called without mapping to a reference assembly. The GBS data are available from the supplemental material deposited on figshare.

### Analysis

Allele frequencies were estimated using allele counts across the relevant group of individuals. Relatedness estimates were calculated using the methods of Dodds *et al.* (2015), including the removal of SNPs with Hardy-Weinberg disequilibrium (observed frequency of the reference allele homozygote minus its expected value) below −0.05. Population structure was visualised by plotting the first two components of a principal components analysis of the genomic relatedness matrix (GRM). Additional calculations were made and used for assigning parentage as described above. Parentage calculations were made twice, firstly using all animals in the dataset, using combined allele frequencies and then separately for each breed, using breed specific allele frequencies. SNPs which had a minor allele frequency (MAF) of zero in the breed-specific set were discarded for that breed.

## Results

### Combined breed analysis

Unless stated otherwise, the relatedness has been used to determine best matching parents. The sequencing and SNP calling resulted in 78,042 SNPs for analysis with a mean read depth of 3.6 and call rate of 72%. The fin plot, which shows apparent Hardy-Weinberg disequilibrium plotted against minor allele frequencies (MAFs), is shown in Figure S2. SNPs with Hardy-Weinberg disequilibrium below −0.05 tended to have high depth and near minimum disequilibrium and may represent reads from duplicated regions. These were removed from further analysis, leaving 77,473 SNPs. The distribution of minor allele frequencies (MAFs) of these SNPs is shown in Figure S3 and shows that a high proportion of SNPs had low MAF. The first two principal components of the GRM are shown in Figure 1. The first component, explains 96.9% of the variation, while the second component explains 0.3%.

**Figure 1.**
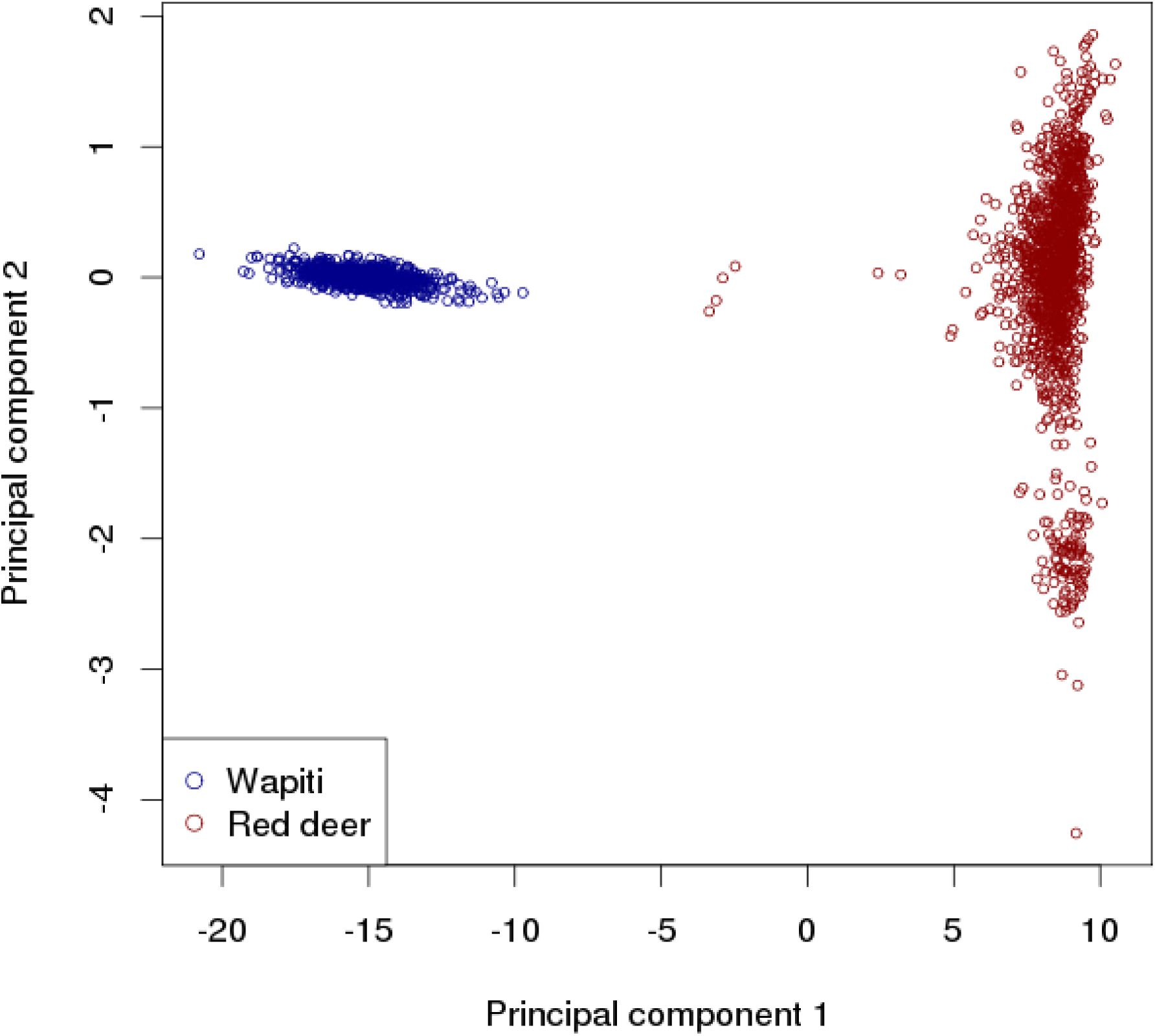
First two principal components from a principal components analysis of the GRM for the combined breed data.

Figure 2A shows the relatedness between each progeny and the best matching sire (i.e. the individual in the sire’s group with the highest relatedness with that progeny). Also plotted is the raw mismatch rate, i.e. the proportion of SNPs whose raw calls are inconsistent with parentage. A plausible relatedness threshold for declaring parentage is a value near 0.5, perhaps a little less than 0.5 to allow for errors in the estimation process. A threshold of 0.4 is shown in Figure 2 (and other relevant plots) as an initial threshold. All but 11 of the progeny have a best matching sire relatedness greater than 0.4, while only two were between 0.4 and 0.5. The other values are considerably higher, averaging 0.61 for Red deer and 0.84 for Wapiti. The raw mismatch rates amongst those above the 0.4 relatedness threshold vary greatly, ranging from 0.011 to 0.065. Raw mismatch rates for the other progeny were all greater than 0.040. Figure 2B shows a similar plot, but with excess mismatch rate (EMM) on the vertical axis. Here there is a clear relationship between relatedness and EMM. All progeny with a best relatedness of at least 0.5 have an EMM below 0.007 while the other progeny have an EMM above 0.019. This suggests an EMM threshold between 0.01 and 0.015, and a relatedness threshold of around 0.5 for declaring parentage, although the EMM threshold combined with any lower relatedness threshold does not change the assignments in these data. Here an EMM threshold of 0.01 is shown in the figures and used for assignments.

**Figure 2.**
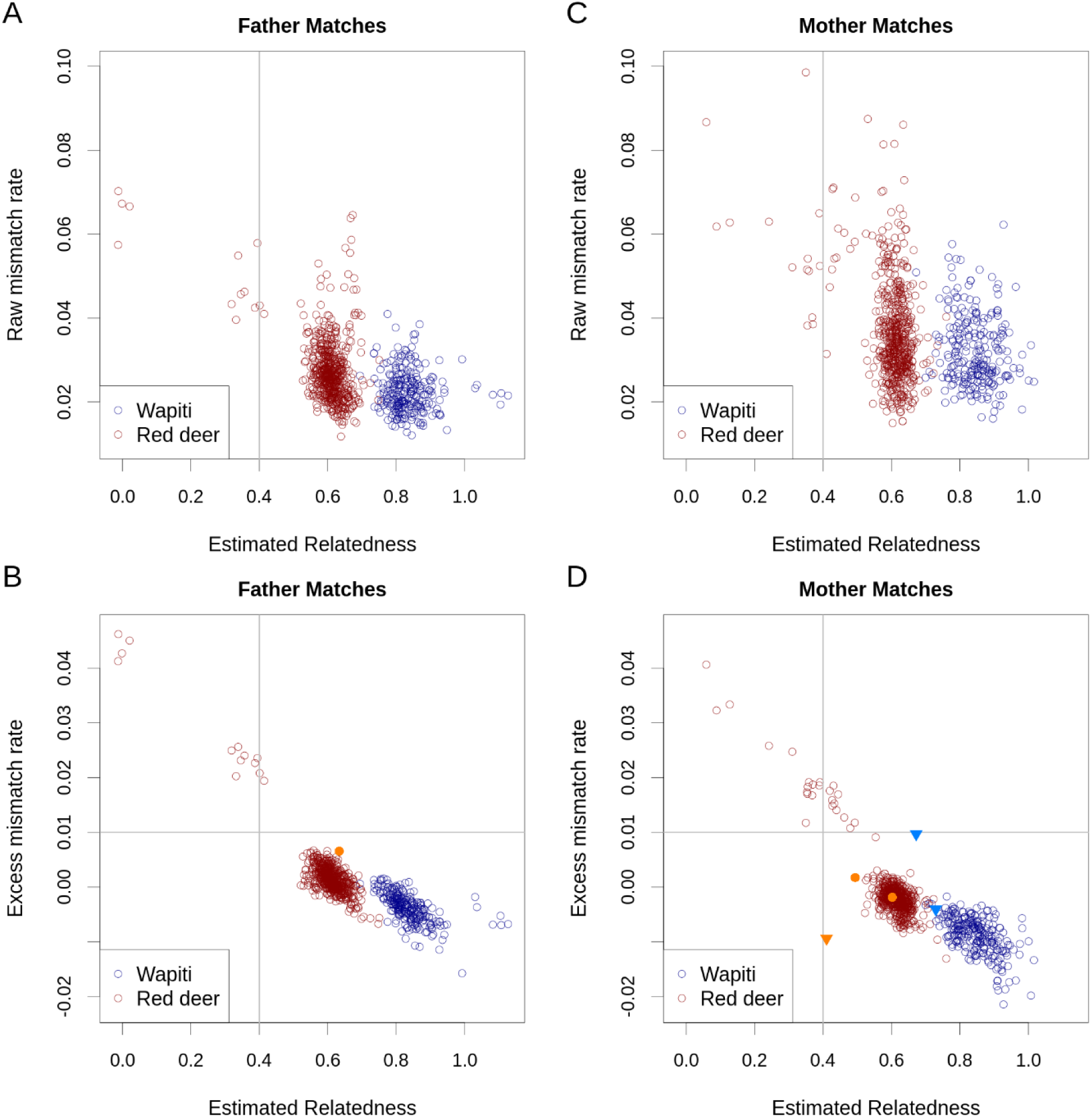
Raw mismatch rate (A, C) or excess mismatch rate (B, D) versus estimated relatedness between the best matching sire (A, B) or dam (C, D) and each progeny for the combined breed analysis. The vertical and horizontal grey lines denote the relatedness and excess mismatch rate thresholds, respectively, for declaring parentage. In B and D, points corresponding to cases where another potential parent has offspring-parent relatedness within 0.05 of plotted value are shown with filled symbols in a brighter shade of their breed colour; those not reaching the 0.99 bootstrap support threshold are shown as triangles.

The corresponding relatedness plots for the best matching dams are shown in Figure 2C (with raw mismatch rates) and Figure 2D (with EMMs). These show a similar pattern to the sire plots. Pairs with an EMM less than 0.009 all had relatedness greater than 0.4 (all but one were greater than 0.49), suggesting relatedness and EMM thresholds of around 0.4 and 0.009, respectively as appropriate threshold for declaring parentage.

One sire assignment and five dam assignments, passing the relatedness (0.4) and EMM (0.01) thresholds, had another potential parent with offspring-parent relatedness within 0.05 of the best matching pair (see Figure S4). If a conservative threshold of 0.99 is used for bootstrap support, three of the dam assignments are rejected, while the other four assignments are retained.

When considering both parent matches together, there were three assignments where the trio EMM exceeded 0.02. It is likely that a higher threshold is appropriate for the trio than pair EMM thresholds, as an error in any of the three individual’s genotypes may generate a mismatch, and so 0.02 is used here as the threshold. For one of these progeny, the second best matching sire and best matching dam gave a low trio EMM (0.006) and this combination passed the other parentage thresholds. Two best sire and dam match trios that passed the relatedness and EMM threshold for each offspring-parent pair had parent relatedness exceeding twice the offspring inbreeding by more than 0.2 (the ‘inbreeding threshold’ being used here to reject a parent pair). One of these was the trio just mentioned (where the second best sire and best dam combination appeared better); the other passed the trio EMM threshold, but had a lower EMM with the best sire and second best dam.

There was only one case where the lowest EMM parent was not one of the two highest related parents. This was for a dam of one of the Red calves. The highest related dam was excluded with the EMM threshold. In this case, the microsatellite-based analysis did not give a dam assignment.

### Separate breed analysis

The combined breed analysis showed quite different relatedness values in the two breeds. In particular, relatedness to the best matching parent for Wapiti is high. The separate breed analysis uses allele frequencies from within each breed, and this may be more appropriate. There were 69,223 SNPs remaining polymorphic within the Wapiti herd and 76,926 in the Red deer herd. Mean SNP depths were similar to the combined analysis (3.8 and 3.3 for Wapiti and Red deer, respectively).

Figure 3 shows the excess mismatch rate and relatedness between each progeny and the best matching sire and dam for the separate breeds analyses. It appears that the thresholds of 0.4 for relatedness and 0.01 for EMM (shown on the plots), or slightly more lenient thresholds, are suitable for declaring parentage. For sire assignment, one Wapiti progeny fails this relatedness threshold (Figure 3A), while 13 Red progeny fail both thresholds (none failing only one threshold, (Figure 3C). For dam assignment, these same thresholds excluded four Wapiti progeny ((Figure 3B; one failing both thresholds, three failing only the relatedness threshold) and 28 Red progeny ((Figure 3D; 24 failing both thresholds, one failing the EMM threshold only and three failing the relatedness threshold only). Most progeny that failed on only one of the thresholds were close to that threshold.

**Figure 3.**
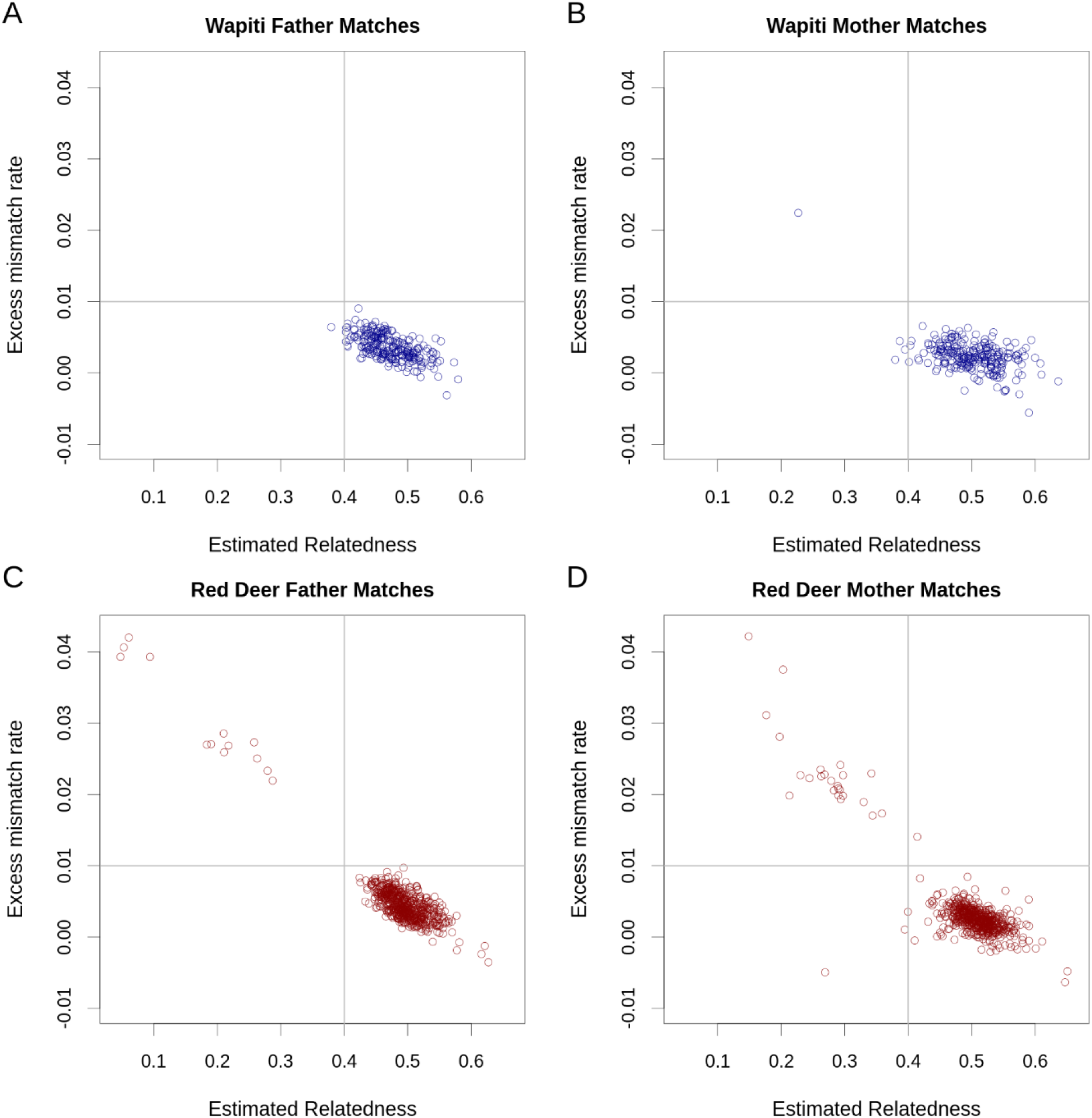
Excess mismatch rate versus estimated relatedness for the best matching sire (A and C) and dam (B and D) with each progeny for the Wapiti (A and B) and Red deer (C and D) analyses.

All these Wapiti assignments passed the trio EMM threshold, but one failed the inbreeding threshold (parent relatedness near zero, but inbreeding just below −0.1). In the Red herd, there were six trios (passing all the single parent thresholds) that failed the trio EMM threshold. One of these also failed the inbreeding threshold. For this progeny the second best sire along with the best dam formed a trio that passed all the thresholds (as was the case for this progeny in the combined breed analysis).

Using the provisional thresholds discussed here, 13 Red deer did not get a sire assignment and 1 Wapiti and 25 Red deer did not get a dam assignment in the combined analysis while 13 Red deer and 1 Wapiti did not get a sire assignment and 4 Wapiti and 28 Red deer did not get a dam assignment in the separate breed analysis. All progeny that had a parent assigned in either of the combined breed or separate breed analysis had the same best matching parent in the other analysis (even if it did not pass the thresholds) except for one Wapiti mother.

### Comparison with microsatellite-based parentage assignments

The separate breed analysis gave relatedness values more in line with that expected and the same thresholds appeared suitable for both breeds. Therefore, results from the separate breed analysis are used for comparison with the microsatellite results.

There were 793 progeny with a sire assignment (after applying all criteria) using GBS; two were assigned a different sire and 791 had the same sire assigned using microsatellites. There were 775 progeny with a dam assignment using GBS. Using microsatellites ten of these were unassigned, two were assigned a different dam and 763 had the same dam assigned.

Eight progeny that passed the single parent thresholds were above the trio EMM (with GBS). Of these, one had a different dam assignment and one had a different sire assignment with microsatellites, compared to the best matching parents with GBS. The one with the different sire assignment also failed the inbreeding test, while the trio with the second best matching sire (the one assigned using the microsatellite test) passed the EMM and inbreeding tests. One additional trio failed the inbreeding threshold but passed the EMM threshold; this trio was assigned by the microsatellite test.

### Alternative allele sampling models

Figure 4 compares raw and expected mismatch rates. It can be seen that the raw mismatch rates are above the expected rates for the majority of trios, including those we have accepted as representing parent-progeny trios (shown with ‘Assign code’ Y). For these trios there is a strong relationship (roughly linear) between raw and expected values, suggesting a systematic reason for the difference. To investigate this further, two alternative allele sampling models (BB and MP) were considered.

**Figure 4.**
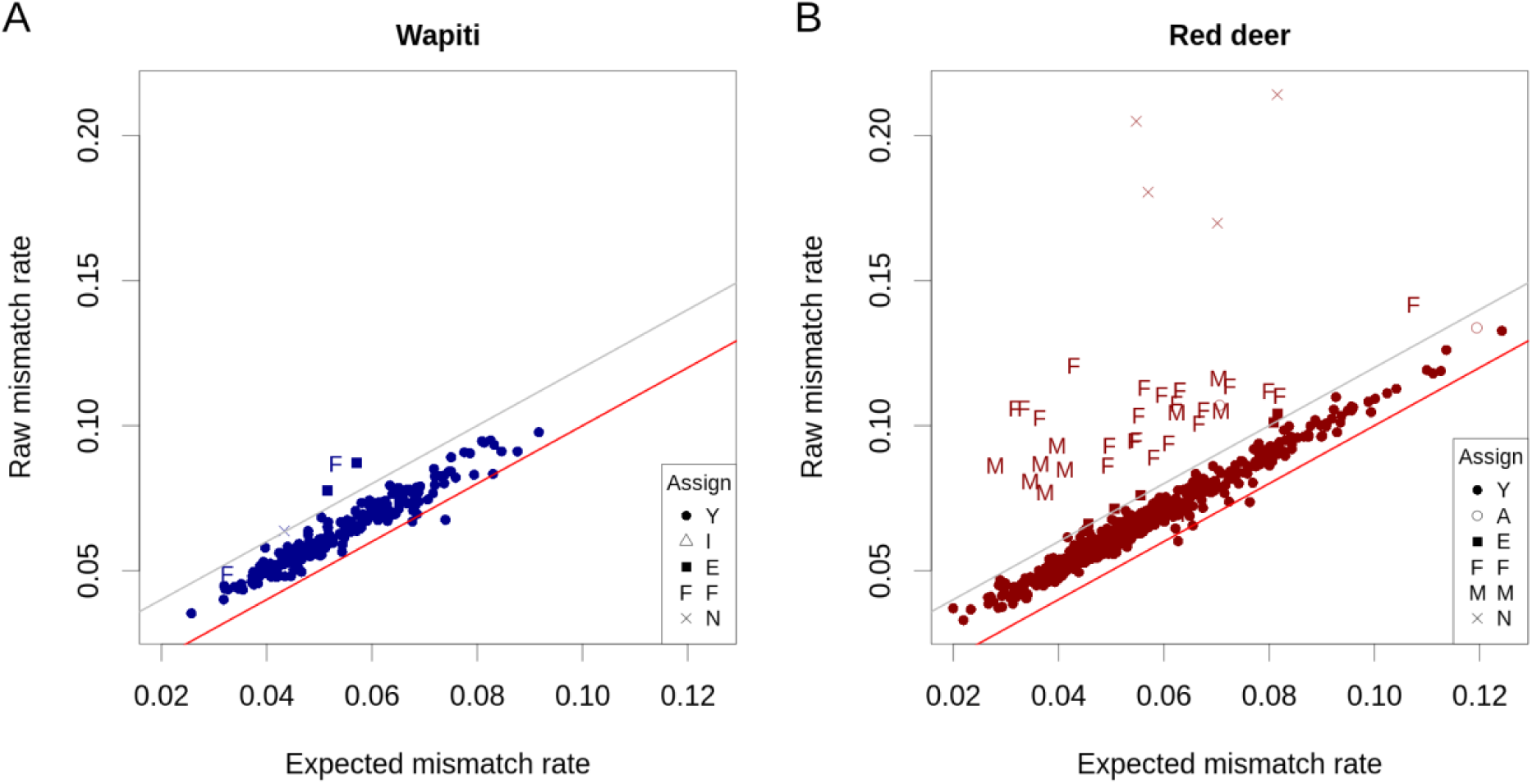
Comparison of raw and expected parent-offspring trio mismatch rates for the Wapiti (A) and Red deer (B) analyses. The red lines show where these are equal. The grey lines show the threshold used for excluding a trio from parentage. Assign codes are Y: assign parentage, A: an alternate parentage has lower EMM, I: fails the inbreeding criterion, E: exclude based on trio EMM, F: assign father only, M: assign mother only, N: do not assign either parent.

The estimated values of β (for the BB model) were 3.96 for the Red herd and 4.61 for the Wapiti herd. The estimated values of *p*′ (for the MP model) were 0.604 for the Red herd and 0.591 for the Wapiti herd. For both herds the BB model gave a lower sums of squares than the MP model. Figure 5 shows raw mismatch rates compared to expected rates calculated with the BB model for the Red herd. All four herd x model combinations are given in Figure S5.

**Figure 5.**
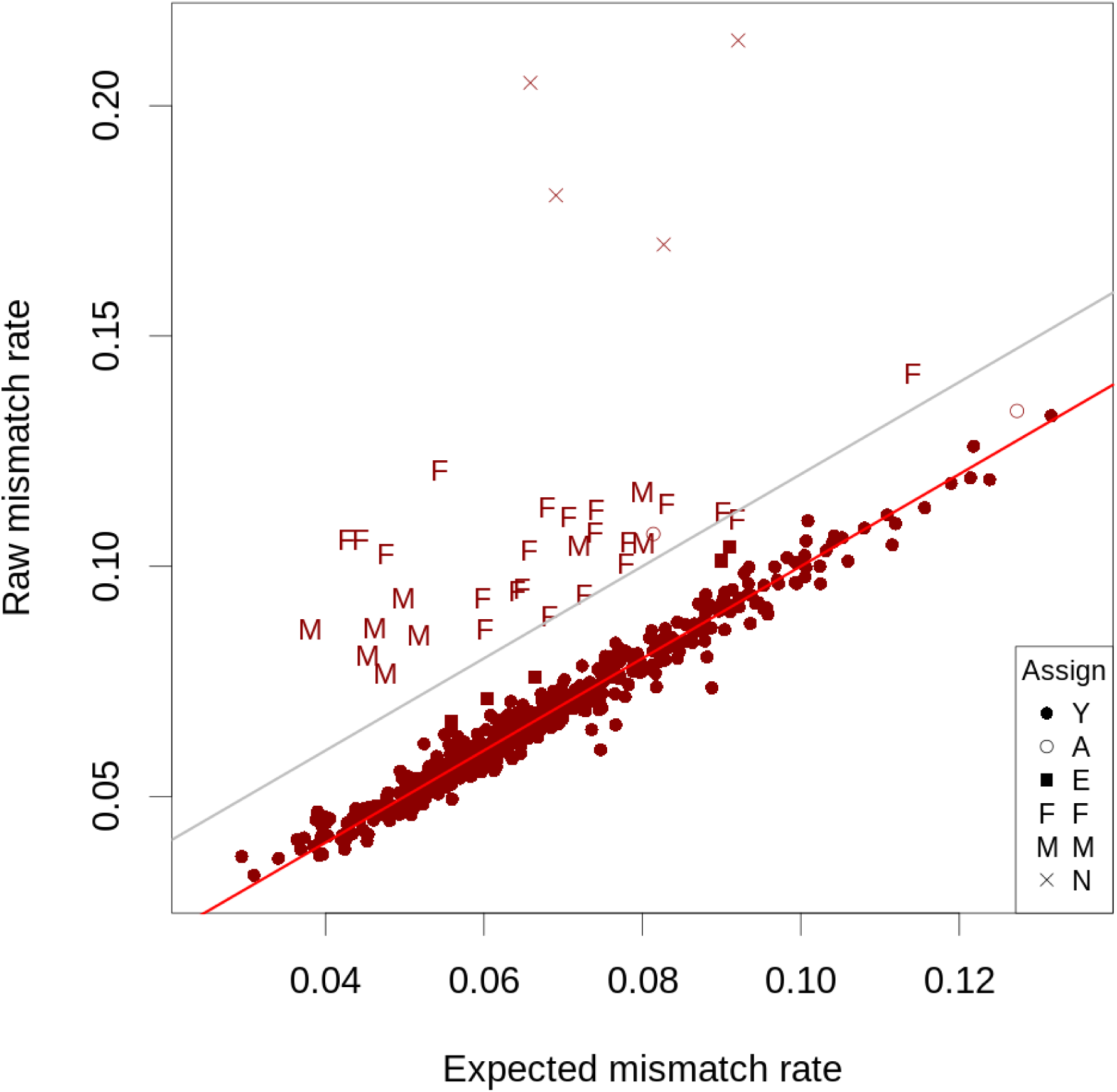
Comparison of raw and expected parent-offspring trio mismatch rates for the Red deer analysis using the BB model. The red line shows where raw and expected rates are equal. The grey line shows the threshold used for excluding a trio from parentage. Assign codes shown are from the analysis using the binomial model; see Figure 4 caption for explanation of these symbols.

## Discussion

We have investigated methods for assigning parentage based on low depth sequencing data. The usual approach to this is to filter genotype calls based on read depth and thereafter assume that genotypes are called with high accuracy. Some software packages (Marshall *et al.* 1998) allow genotype calls with errors, using a supplied error rate that is constant across all genotypes. However here the actual error rate (due to allele sampling) is dependent on the genotype call and its read depth.

Modelling the sequencing process, analogous to that used by Dodds *et al.* (2015) for relatedness estimation, Bilton *et al.* (2018b) for linkage analysis and Bilton *et al.* (2018a) for linkage disequilibrium estimation, allows depth-dependent errors. Recently, Whalen *et al.* (2019) have adopted this approach to develop likelihood-based methods for relationship classification (including parentage). Here, we have considered the use of relatedness estimates using the KGD method of Dodds *et al.* (2015) and expected mismatch rates. Both these statistics model the probability of observing both alleles for a heterozygote in a depth-dependent way assuming known allele frequencies. Excess mismatch rates (EMM) are then calculated as the difference between raw (assuming the genotype is given by the alleles observed) and expected rates. The EMM is similar to mismatch rates used in exclusion-based parentage assignment methods that allow for some genotyping error. The EMM could be calculated after filtering on read depth, but as it accounts for depth, it allows all SNPs (not filtered out for other reasons) to be used. In practice, it may suffice to calculate the EMM only for the leading candidate parents, based on relatedness estimates. This leads to a computationally efficient approach, as estimated relatedness between pairs can be calculated faster than the EMM (approximately 10x faster using current software).

The analysis of the deer data showed that the raw mismatch rate was not a reliable indicator of parentage – there was considerable overlap of these values amongst pairs that were either consistent or not with parentage, based on relatedness estimates. In contrast, it was possible to find relatedness and EMM thresholds for declaring parentage that were almost fully concordant, with only one case where the two methods gave different conclusions. This case did have a low EMM (−0.005) but with a low relatedness (0.27) for the best mother match in the Red deer analysis. The same case was less of an outlier in the combined breed analysis (EMM of −0.009, relatedness of 0.41), showing that the results are dependent on the allele frequencies used.

The choice of allele frequencies used influences the measures to determine parentage. The theory assumes that allele frequencies for the ‘base population’ are known. The two breeds examined in this study were clearly separated by the 1^st^ principal component (a few intermediate Red deer probably had some Wapiti ancestry). For the combined breed analysis we used estimates based on the total allele counts of all animals genotyped in the study. This resulted in high relatedness estimates within herds (for example, as demonstrated by average relatedness between progeny and best matching sire of 0.84 and 0.60 for the Wapiti and Red herds, respectively). The allele frequencies used may not be a very good representation of the allele frequencies before breed differentiation, due to genetic drift within these two breeds. For example, there are negative estimates of relatedness. For the within herd analysis, the relatedness values dropped, reflecting that the allele frequencies relate to the genetic variation within each herd. The within herd estimates between progeny and best matching parent are more in the range that would be expected for parent-offspring pairs with unrelated parents. Therefore, choosing an appropriate base population for allele frequency estimation may help in choosing thresholds that seem sensible. A within-breed analysis would normally be preferred. Despite the differences in the relatedness estimates using the within breed compared to combined breed allele frequencies, the two analyses were generally consistent in terms of parent assignment. Using a set of provisional thresholds, any time an assignment was made in both analyses it was to the same parent. This would still be the case if the relatedness threshold in the separate breed analyses was dropped to 0.35 (7 more assignments). There was one case of a combined analysis mother assignment which had a different best matching mother in the Wapiti analysis. For this case the EMM was 0.0094, close to the threshold (0.01). A slightly lower EMM threshold may be more appropriate in the combined breed analysis (Figure 2). Even amongst cases not assigned by either analysis, most (31 out of 39) had the same best matching parents. These observations suggest that if appropriate thresholds are chosen, the assignments may be reasonably consistent with respect to the use of different allele frequencies. The method requires appropriate thresholds which can usually be determined when there are sufficient numbers of progeny, similar to what Moore *et al.* (2019) found using genomic relatedness from SNP chip data.

The estimated inbreeding in the offspring plus half that of the putative parent could be subtracted from the parent-offspring relatedness values to adjust for inbreeding. This could change which possible parent has the highest relatedness match. It may bring the values closer to the value of 0.5, as expected for non-inbreds. However, estimated inbreeding is generally less precise than between individual relatedness estimates (Dodds *et al.* 2015), especially with low depth sequencing, so the adjusted relatedness would have more variability than the original estimate. In addition, breeding programmes usually aim to keep levels of inbreeding low and so there may be little variation in inbreeding relative to the assumed base population. In the combined breed analysis, where the population is genetically diverse and subdivided, using the adjusted relatedness approach gave best matching relatedness values mostly between 0.3 and 0.5 (data not shown) Although the approach reduced the variation in these values, it appears to have over-corrected the relatedness estimates in this case.

The (within breed) GBS-based assignments made here were largely consistent with assignments based on a microsatellite parentage panel. There were four cases of a different parent assignment. Two were the only ones assigned to a particular sire using microsatellites while this sire had no progeny assigned with GBS. Different DNA samples from the sire were used for microsatellites and GBS, so it is possible there was a sample mix-up. The other difference was for a mother assignment, with no clear reason for the difference; a sample mis-identification is possible. The use of trio information (trio EMM and the inbreeding test) were found to be useful in reducing the number of mismatches between the methods, and such information should be used where possible. These tests help prevent false assignments where a true parent is missing, but a close relative of the other parent of the offspring meets the relatedness and single parent EMM thresholds (e.g. helps avoid assigning a paternal aunt as the mother).

The EMM tended to be greater than zero, even for cases which appeared to be correct matches. A possible reason for this is that the alleles at a locus are not sampled randomly but there tends to be clustering in the alleles recruited for sequencing. This could be due, for example, to PCR artefacts such as “stacking” (Andrews *et al.* 2016) or suboptimal ratios of reagents (Ott *et al.* 2017). Two alternative allele sampling models were proposed and applied, the beta-binomial model and a model where the first allele is sampled at random, but subsequently the same allele is more likely to be sequenced than the other allele. Both models improved the EMM values by similar amounts, such that the EMM tended to be close to zero for what appeared to be correct matches, with the beta-binomial model giving a better fit. The use of either of these models would allow tighter EMM thresholds and may lead to more accurate parentage assignments. These models could also be applied in other GBS analyses, such as the estimation of inbreeding (Dodds *et al.* 2015), linkage (Bilton *et al.* 2018b), linkage disequilibrium (Bilton *et al.* 2018a), calculating genotype likelihoods for downstream analyses (Korneliussen *et al.* 2014) or predicting gender (Bilton *et al.* 2019). More work is required to evaluate whether these differences are important, whether the alternate models are significantly better than the random sampling model or whether there are other models (for example, one which allows sequencing error) which are more realistic and provide a better fit.

## Conclusions

The developments described here provide methods and guidelines for assigning parentage from sequencing data. These methods have been developed with reduced representational low depth sequencing in mind. This allows sequencing costs to be as low as possible making it a possibility for parentage analysis, perhaps in combination with other genomic analyses. The sequencing approach allows genomic resources to be developed with low start-up costs compared to other approaches. The parentage assignment process requires some choice of appropriate thresholds, but these are usually clear-cut. A possible exception is where the population is genetically diverse with distinct subgroups, and there are different possible approaches to calculating allele frequencies to use in the process. When applied to a deer dataset, the parentage assignment was largely consistent with assignments made using a microsatellite panel; the few differences seen may have been due to sample tracking errors. Therefore, GBS or other sequencing-based methods can be used for parentage assignments, increasing the utility of these sequencing methods.

## Acknowledgements

This project was supported by the Ministry of Business, Innovation and Employment via its funding of the “Genomics for Production & Security in a Biological Economy” programme (Contract ID C10X1306) and FarmIQ (Ministry for Primary Industries’ Primary Growth Partnership fund) – FIQ Systems – Plate to Pasture (Reference: PGP06-09020). We thank Focus Genetics (New Zealand) and Pāmu (formerly Landcorp Farming Ltd) (New Zealand) for the use of their animals and genotypes for testing the proposed methods.

